# Development of a dual chemical probe for the USP16 and HDAC6 zinc-finger ubiquitin-binding domain

**DOI:** 10.1101/2025.10.23.684218

**Authors:** Madhushika Silva, Mandeep K. Mann, Bijan Mirabi, Magdalena M. Szewczyk, Joachim Loup, Aurelien Dupeux, Renu Chandrasekaran, Jonathan Bajohr, Cheryl H. Arrowsmith, Matthieu Schapira, Mark Lautens, Dalia Barsyte-Lovejoy, Rachel J. Harding, Vijayaratnam Santhakumar

## Abstract

Ubiquitin-specific peptidase 16 (USP16) is a deubiquitinase that specifically cleaves ubiquitin from histone H2A, and modulates gene expression, cell cycle regulation, and various other cellular processes. The USP16 zinc-finger ubiquitin-binding domain (UBD) binds the free C-terminal end of both ubiquitin and ISG15, two major signaling proteins that mediate many biological pathways. Because the precise function of USP16-UBD and its interactions remains unclear, a small molecule antagonist targeting the USP16-UBD could enable cellular studies to elucidate its biological role. Here we report SGC-UBD1031 (**15**), a chemical probe targeting USP16-UBD with similar *in vitro* binding profiles to HDAC6-UBD and selectivity over nine other UBDs. In cellular assays, **15** disrupts the interaction between the C-terminus of ISG15 and USP1-UBD, as well as the interaction between ISG15 and HDAC6 UBD, at a concentration of 1 μM. The corresponding enantiomer SGC-UBD1031N (**16**), does not interfere with these interactions, even at concentrations as high as 30 μM, and thus serves as a negative control.

**Table of Contents graphic:** 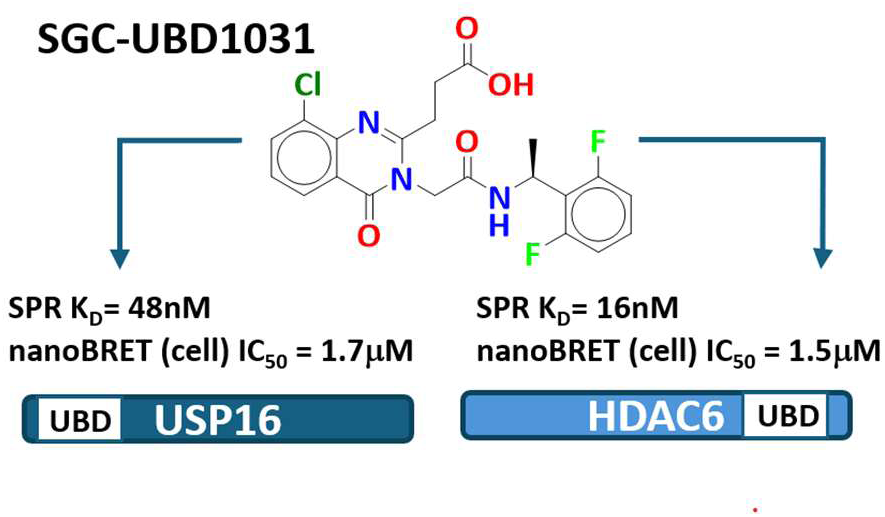

## INTRODUCTION

Ubiquitin-specific peptidase 16 (USP16) is one of approximately 100 deubiquitinases (DUBs) encoded in the human genome. Comprising an 823 amino acid molecule, USP16 consists of an N-terminal UBP-type zinc-finger domain (UBD) and a catalytic domain (aa. 193–823) that is bisected by a disordered region (aa. 393–627) ^1^. USP16 deubiquitinates histone H2A at Lys-119 ^2^ and its increased expression has been associated with a wide range of cellular and pathological abnormalities due to enhanced H2A deubiquitination. Studies have shown that embryonic stem cell pluripotency depends on USP16 mediated removal of ubiquitin from histone H2A, a process that silences lineage-specific genes and prevents premature differentiation ^3^. Conditional knock out of USP16 in murine bone marrow significantly elevates global ubiquitinated H2A levels, leading to lethality ^4^ while its depletion in HeLa cells slows growth by disrupting the mitotic phase of the cell cycle ^5,6^. Aberrant USP16 activity is implicated in cancer progression, where USP16 overexpression is associated with increased proliferation and tumorigenesis ^7–9^. USP16 is also known to play a critical role in regulating the DNA damage response by removing ubiquitin marks to restore transcription and resolve repair signals. However, its misregulation, whether through overexpression or abnormal nuclear accumulation, disrupts proper DNA repair processes^10–12^. Additionally, USP16 is also implicated in other pathologies, including autoimmune diseases ^13,14^ Down’s syndrome ^15–17^ and male infertility ^18^.

In our previous work ^19^, we reported the development of the first potent and selective small molecule chemical probe targeting the HDAC6-UBD, which disrupts the interaction between the C-terminal ubiquitin RLRGG peptide and HDAC6, with selectivity over the closely related USP16. Here, we describe the exploration of the structure-activity relationship (SAR) of this chemical series to improve USP16-UBD binding activity, and characterization of SGC-UBD1031 (**15**), a potent antagonist targeting both USP16-UBD and HDAC6-UBD. This chemical probe exhibits comparable activity against both UBDs, demonstrating efficacy in cells at 1 μM, high selectivity over other closely related UBDs and no detectable toxicity. We identified the corresponding *R-*enantiomer, SGC-UBD1031N, as a negative control, with >30-fold reduced activity for both USP16-UBD and HDAC6-UBD. Binding parameters for SGC-UBD1031 and SGC-UBD1031N were determined using surface plasmon resonance (SPR) and isothermal titration calorimetry (ITC). Our results show that **15** effectively disrupts the interaction of ISG15 with both USP16-UBD and HDAC6-UBD in cells at 1 μM, while the negative control **16** shows no activity across all concentrations tested. These tool compounds will facilitate the investigation of USP16 function when used alongside the previously reported HDAC6 UBD-selective probe.

## RESULTS

### STRUCTURE−ACTIVITY RELATIONSHIP

We previously reported compound **1**, a potent, selective, and cell-active chemical probe targeting the HDAC6-UBD. To further improve the cellular activity of this HDAC6-UBD probe, we introduced a methyl group at the benzylic position (**3**), which resulted in a threefold improvement in binding affinity to USP16-UBD, from 1.3 μM to 0.44 μM, while maintaining its binding to HDAC6-UBD, as measured by SPR. In contrast, the introduction of an N-methyl group in the negative control (**2**) led to a >300-fold reduction in HDAC6 binding activity, with minimal effect on USP16 binding. This observation prompted us to combine these two modifications—the benzylic methyl group to enhance USP16-UBD binding and the N-methyl group to reduce HDAC6-UBD binding (**4**) —in an attempt to develop a USP16-UBD-selective probe. However, this approach resulted in the complete loss of binding activity for both USP16-UBD and HDAC6-UBD (Figure 1).

**Figure 1.**
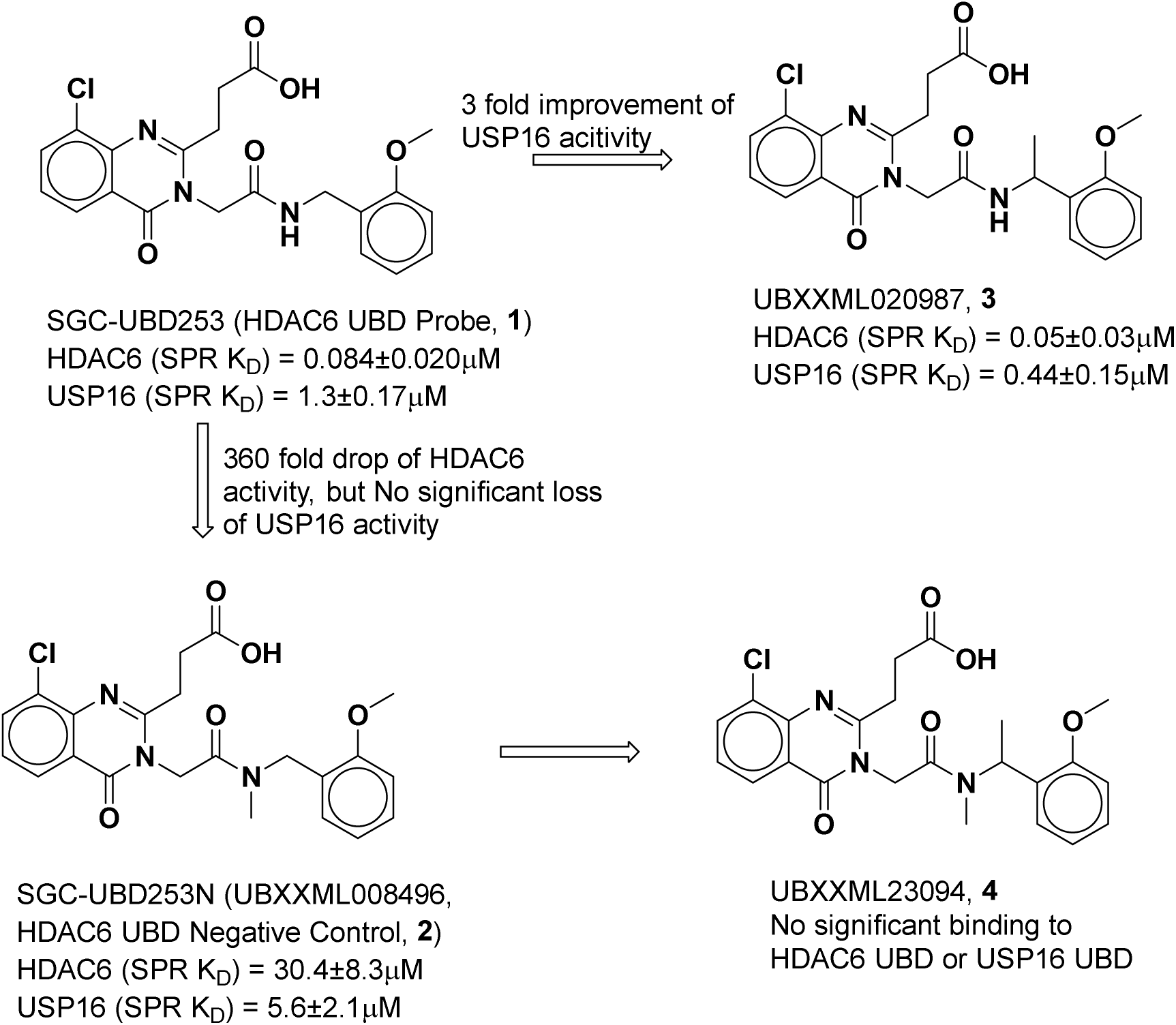
USP16 and HDAC6 activity of the methylated analogues of the HDAC6 probe (1)

To further guide structure-based optimization, we attempted to co-crystallize compound **3** with USP16-UBD; however, all attempts were unsuccessful. Nevertheless, the superimposition of the co-crystal structure of HDAC6-UBD with its probe (**1**) (PDB 8G45) and the reported NMR structure of apo USP16-UBD (PDB 2I50) ^20^ revealed several key differences (Figure 2). In USP16-UBD, the arginine R67 is positioned away from the compound binding site, in contrast to the corresponding arginine residue, R115, in HDAC6, that forms a π-interaction with the quinazolinone core and a salt bridge with the carboxyl group of **1**. Notably, this arginine in HDAC6 is highly flexible, adopting different orientations in both the apo structure (PDB 3C5K) and the co-crystal with the peptide (PDB 3GV4), as well as in the co-crystal structure with the HDAC6 probe compound (PDB 8G45). This flexibility suggests that a similar conformational shift may occur for R67 in USP16 upon compound binding. Additionally, the E105 in USP16 for the corresponding Y1189 in HDAC6 in the pocket may account for the apparent tolerance of N-methyl substitutions by USP16, which is not tolerated by HDAC6. Despite this, the NH group is still expected to form a hydrogen bond with D105 in USP16, which is crucial for binding. Key differences also exist in the residues lining the side pocket, where USP16 contains N94, D93, and S53, in contrast to D1178, I1177, and E1141, respectively at the corresponding positions in HDAC6. To leverage these differences in the side pocket for achieving USP16 selectivity, we explored a range of amide substitutions replacing the methoxy benzyl group in **1**. The results are summarized in Table 1. These compounds were synthesized following the same general procedures used for the HDAC6 probe compound (SGC-UBD253).

**Figure 2.**
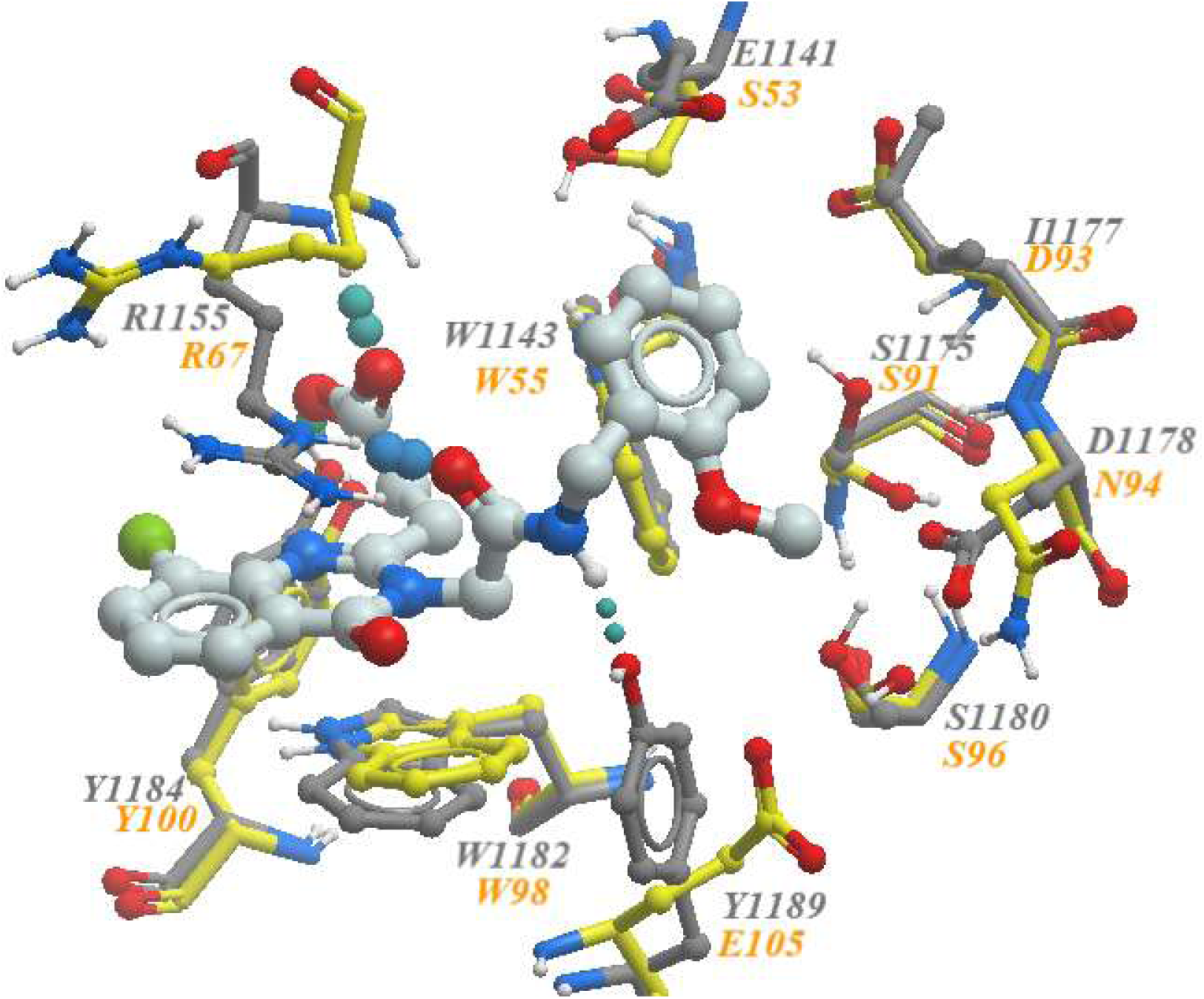
Co-crystal structure of **1** in complex with HDAC6-UBD (**1** shown in pale blue and HDAC6-UBD sidechains in grey, (PDB ID 8G45) superimposed with the NMR structure of USP16-UBD (key residues shown in yellow, PDB ID 2I50) showing key differences in the adjacent pocket.

**Table 1.**
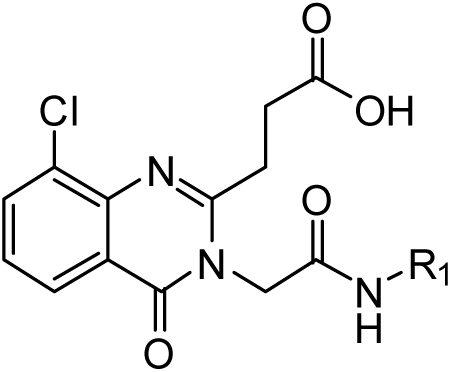
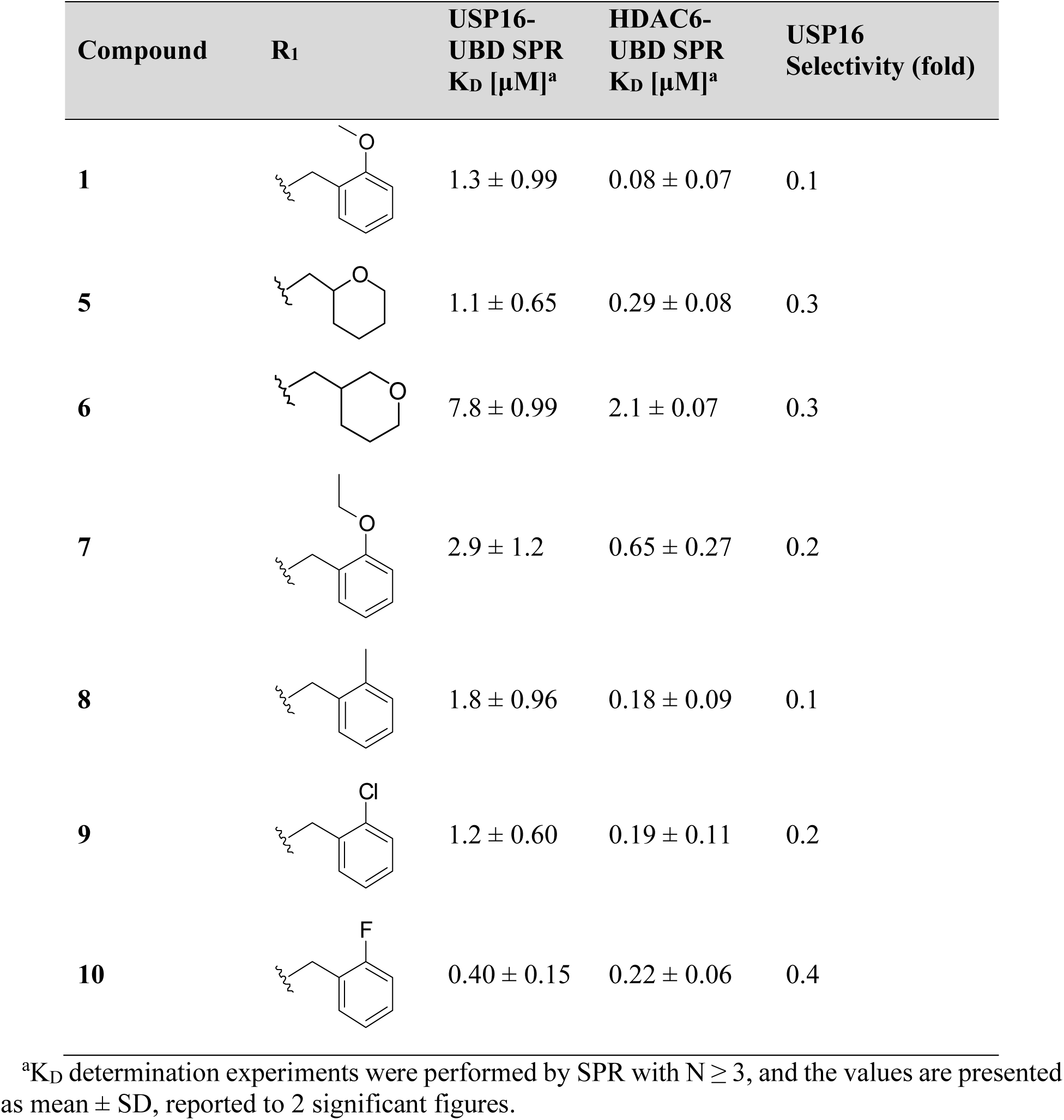
USP16 and HDAC6 activity of amide substitutions (1, 5-10)

| Compound | R<sub>1</sub> | USP16-UBD SPR<br>K<sub>D</sub> [μM]<sup>a</sup> | HDAC6-UBD SPR<br>K<sub>D</sub> [μM]<sup>a</sup> | USP16<br>Selectivity (fold) |
| --- | --- | --- | --- | --- |
| 1 |  | 1.3 ± 0.99 | 0.08 ± 0.07 | 0.1 |
| 5 |  | 1.1 ± 0.65 | 0.29 ± 0.08 | 0.3 |
| 6 |  | 7.8 ± 0.99 | 2.1 ± 0.07 | 0.3 |
| 7 |  | 2.9 ± 1.2 | 0.65 ± 0.27 | 0.2 |
| 8 |  | 1.8 ± 0.96 | 0.18 ± 0.09 | 0.1 |
| 9 |  | 1.2 ± 0.60 | 0.19 ± 0.11 | 0.2 |
| 10 |  | 0.40 ± 0.15 | 0.22 ± 0.06 | 0.4 |

<sup>a</sup>K<sub>D</sub> determination experiments were performed by SPR with N ≥ 3, and the values are presented as mean ± SD, reported to 2 significant figures.

Compared to **1**, the corresponding 2-tetrahydropyran derivative (**5**) exhibited similar USP16 binding activity but a 4-fold decrease in HDAC6 binding. In contrast, the 3-tetrahydropyran derivative (**6**) demonstrated a 6-fold reduction in USP16 binding and a 26-fold reduction in HDAC6 binding. These results suggest that the hydrogen bond between the oxygen atom of compound **1** and Y1189 in HDAC6 plays a crucial role in HDAC6 binding, although it is not critical for USP16 binding, as the oxygen atom cannot form a hydrogen bond with the corresponding residue, E105, in USP16. The corresponding ethoxy analogs (**7**) of compound **1** did not improve USP16 activity or selectivity against HDAC6. However, both the 2-methyl (**8**) and 2-chloro (**9**) analogs retained USP16 binding activity and similar selectivity profiles. Interestingly, only the 2-fluoro analog (**10**) showed a 3-fold improvement in USP16 binding, along with a similar fold decrease in HDAC6 binding, resulting in an overall improvement in USP16 selectivity but still maintaining potent HDAC6 activity.

In addition to the compounds **5–10** summarized in table 1, we also tested several variations of the amide group and compounds with no or other substitution at the quinazoline-4-one ring, which had been synthesized previously (SI, Table 1) in our previous work. However, none of these compounds exhibited significant improvement of USP16 activity. Despite testing a range of variations, only compounds with weak USP16 activity demonstrated moderate selectivity (up to 10-fold) for USP16. Given that we have already reported an HDAC6-selective probe, we chose to focus on developing a dual probe that targets both USP16 and HDAC6 as it would still be a valuable tool for investigating the function of USP16 when used in parallel with the HDAC6-selective probe.

As initially observed with compound **3**, the introduction of a methyl group at the benzylic position of **10** (**12**) resulted in enhanced binding to both HDAC6 and USP16. Similarly, methylation of the N-H group in compounds **10** and **12** to form the *N*-Me derivatives (**11** and **13**, respectively) led to a significant loss of HDAC6 activity and a moderate, approximately 5-10-fold reduction in USP16 activity. However, the addition of a second fluorine atom at the other ortho position (**14**) further improved USP16 binding, reducing the K_D_ to below 100 nM, thus meeting the *in vitro* binding criteria for a chemical probe (Figure 3).

**Figure 3.**
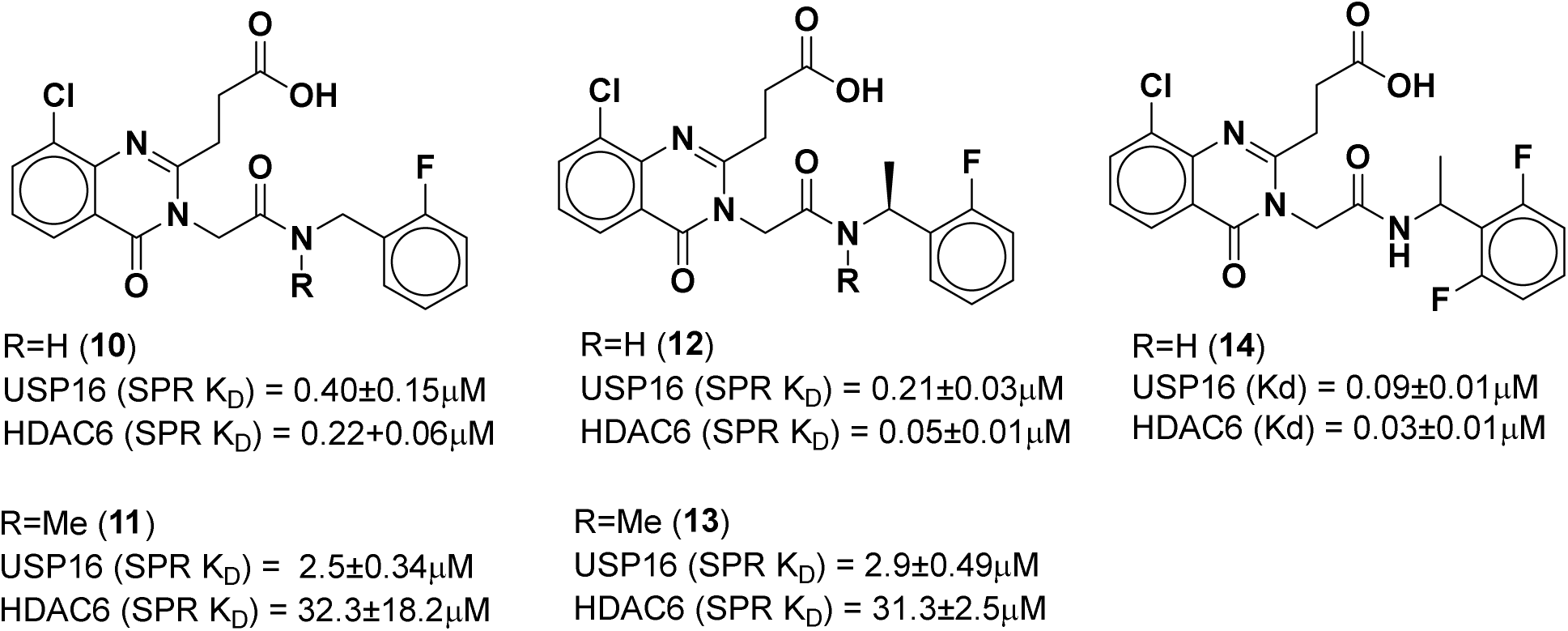
Effect of *N*-Me, α-methylbenzyl and ortho fluorophenyl substitutions on USP16 and HDAC6 binding.

### IN VITRO CHARACTERIZATION OF 15 and 16

We synthesized both enantiomers of **14**, the *S*-enantiomer **15** binds potently to USP16-UBD with K_D_ values of 0.048 μM and 0.05 μM as determined by SPR and ITC, respectively, and binds to HDAC6-UBD with K_D_ values of 0.016 μM as determined by SPR (Figure 4). In contrast, the *R*-enantiomer **16** binds only weakly to USP16-UBD with a K_D_ of 1.7 μM and to HDAC6-UBD with a K_D_ of >5 μM as determined by SPR. The *S*-enantiomer **15** (SGC-UBD1031) was selected as a probe candidate and the corresponding *R*-enantiomer **16** (SGC-UBD1031N), which is > 30-fold less active at both USP16 and HDAC6 was chosen as a negative control candidate and profiled further.

**Figure 4.**
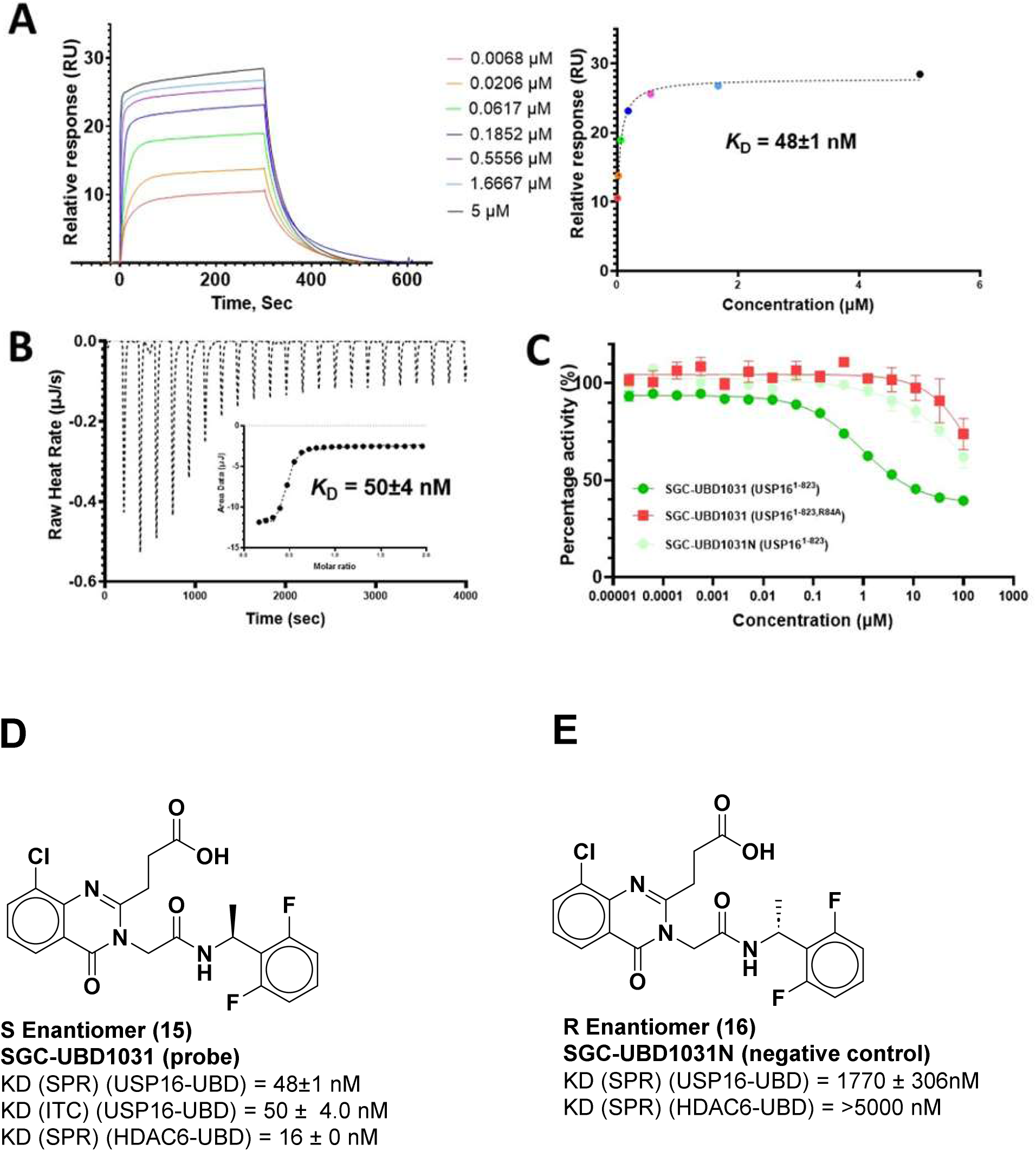
*In vitro* characterization of SGC-UBD1031. A) (left)-SPR sensorgram showing the dose-dependent titration of SGC-UBD1031 binding to USP16-UBD, and (right)-response vs. concentration plot using a steady state response. B) ITC thermogram generated for the injection of SGC-UBD1031 and (inset) corresponding fitted curve used to determine KD values for the interactions. C) Effect of SGC-UBD1031 (dark green) and SGC-UBD1031N (pale green) on DUB activity of USP16-FL, and the control run with - catalytic dead mutant USP16-FL, R84A (light green). D) SGC-UBD1031 (**15**) and E) SGC-UBD1031 (**16**) n=3 for each experiment.

Both probe and negative control were tested to assess possible inhibition of the deubiquitinase activity of USP16. In addition to the full-length USP16(1-823) (USP16-FL), the catalytically dead mutant USP16-FL R84A was also used to investigate the effects of SGC-UBD1031 and SGC-UBD1031N. The probe demonstrates up to 60% inhibition of DUB activity in a dose-dependent manner with IC_50_ 10.6 ± 1.5 µM of USP16-FL (Figure 4) while the negative control shows no inhibition of DUB activity, behaving similarly to the catalytically dead mutant USP16-FL R84A. SGC-UBD1031 demonstrates cooperative inhibition of DUB activity of USP16 via binding to the UBD.

Next, we tested the selectivity of SGC-UBD1031 and SGC-UBD1031N for USP16-UBD against a panel of UBD proteins by SPR, which were selected to represent at least one member from the majority of the branches from the dendrogram derived from the alignment of Zf-UBD pocket residues (Figure 5). These probe and negative control candidates were at least 100-fold selective for USP16 and HDAC6 over 7 other representative UBD proteins tested (Table 2).

**Figure 5:**
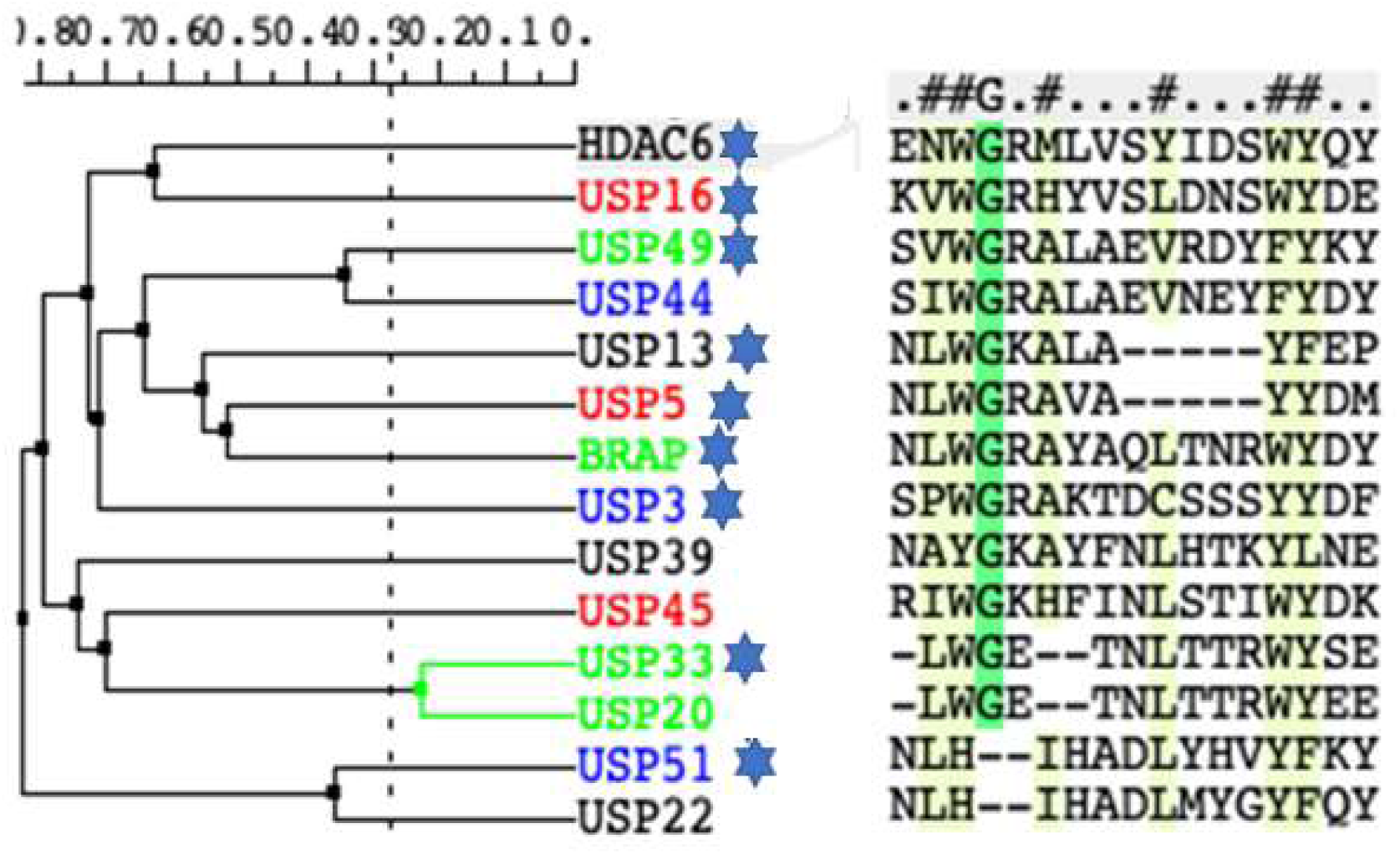
Dendrogram derived from alignment of Zf-UBD pocket residues

**Table 2.** Selectivity of the probe and negative control against a panel of UBD-containing proteins.

| **Protein** | **Probe K<sub>D</sub> (nM)**<br>**SGC-UBD1031** | **Fold selectivity**<br>**compared to**<br>**USP16/HDAC6** | **Negative Control K<sub>D</sub> (nM)**<br>**SGC-UBD1031N** |
| --- | --- | --- | --- |
| USP3 | >10,000 | >100 | 8700 ± 1800 |
| USP5 | 2600 ± 1400 | >100 | >10,000 |
| USP13 | >10,000 | >100 | >10,000 |
| USP16 | 48 ± 1 | - | 1770 ± 300 |
| USP33 | >10,000 | >100 | >10,000 |
| USP49 | 7500 ± 870 | >100 | 6700 ± 2100 |
| USP51 | >10,000 | >100 | >10,000 |
| BRAP | >10,000 | >100 | >10,000 |
| HDAC6 | 16 ± 0 | - | >5000 |

### CHARACTERIZATION OF THE PROBE AND THE NEGATIVE CONTROLS IN CELLS

To directly assess SGC-UBD1031 cellular target engagement in live cells, we utilized the previously described nano-bioluminescence resonance energy transfer (NanoBRET) protein-protein interaction assay (Figure 6A). This assay measures energy transfer from a bioluminescent protein donor (NanoLuc-tagged HDAC6 or USP16) to a protein acceptor (fluorescently labelled HaloTag-tagged ISG15). In HEK293T cells, SGC-UBD1031, but not its negative control enantiomer SGC-UBD1031N, significantly decreased the interaction between HDAC6 or USP16 and ISG15, exhibiting similar potency with IC_50_ values of 1.5 ± 0.2 and 1.7 ± 0.2 μM, respectively (Figure 6B and C). To test the toxicity of both the probe and the negative control compounds, we treated HEK293T and HCT116 cells for 5 days with each compound at 30 μM concentration and monitored cell confluency with Incucyte microscope. Neither of the compounds had any effect on cell growth in either cell line.

**Figure 6.**
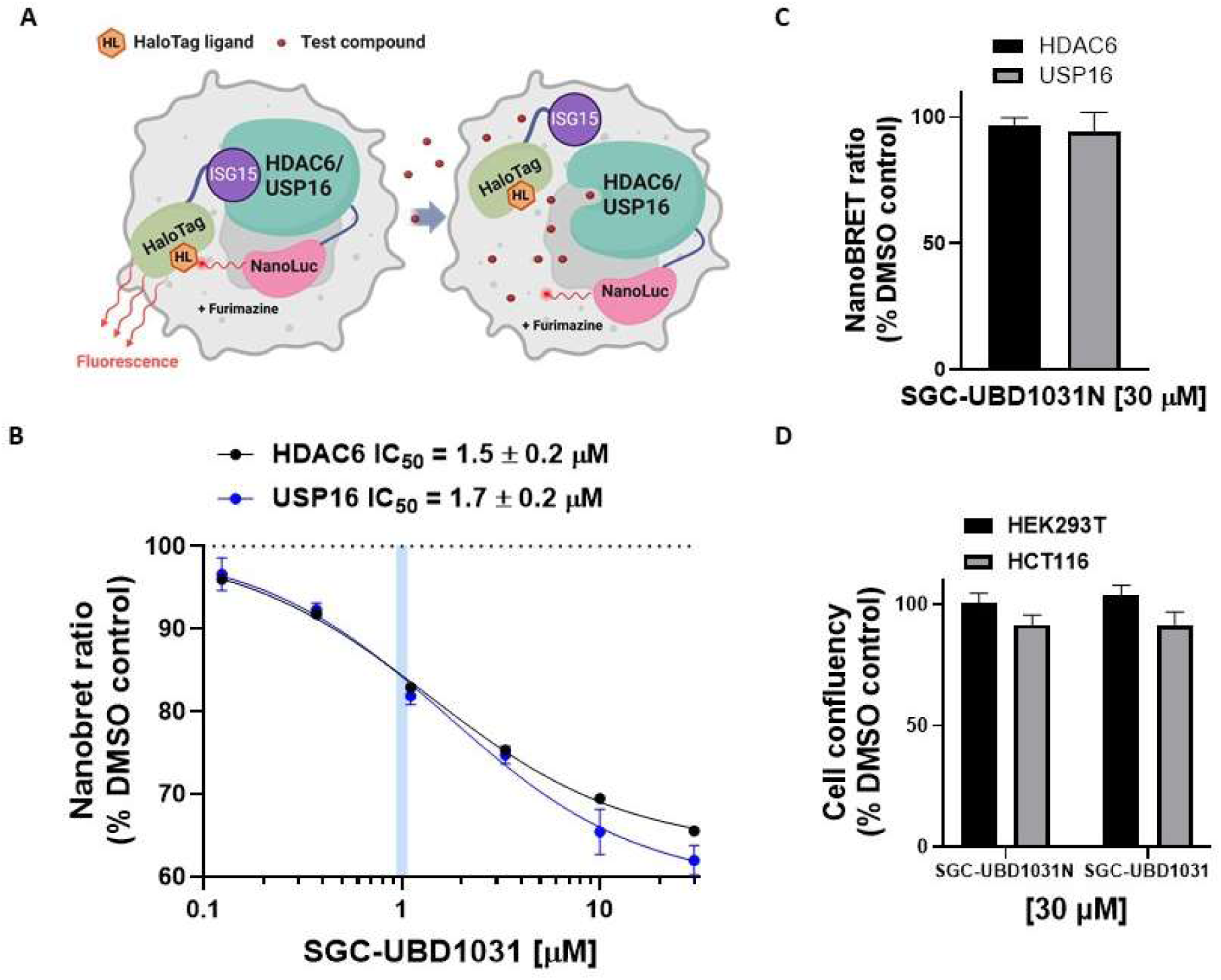
SGC-UBD1031 decreases the interaction between ISG15 and the UBD domain of USP16 and HDAC6 with similar potency. A) Schematic of NanoBRET assay, which measures the BRET between NanoLuc-tagged HDAC6 or USP16 (donor) and HaloTag-tagged ISG15 (acceptor) in the presence of the HaloTag NanoBRET 618 Ligand (HL). Created in BioRender. Barsyte-lovejoy, D. (2025) https://BioRender.com/ mi0rg73. B) SGC-UBD1031 inhibited the interaction of HDAC6 or USP16 and ISG15 in HEK293T cells in a dose-dependent manner (MEAN+/-SEM N=8, 2 biological replicates with for replicates each). C) SGC-UBD1031N did not inhibit the interaction of HDAC6 or USP16 and ISG15 in HEK293T at 30 μM concentration (MEAN+/-SEM n=3, technical replicates). D) SGC-UBD1031 and SGC-UBD1031N did not affect the cell growth of HEK293T and HCT116 cells at 30 μM. Cells were treated with compounds and confluency was monitored for 5 days (MEAN+/-SEM n=3 technical replicates).

## CONCLUSIONS

We report the identification of a dual USP16/HDAC6-UBD chemical probe (**15**) (SGC-UBD1031) and its negative control (**16**) (SGC-UBD1031N), which we thoroughly characterized with *in vitro* biophysical assays and functional cellular target engagement assays. SGC-UBD1031 is a potent antagonist of both USP16-UBD and HDAC6-UBD, active in cells against both targets, and shows good selectivity against other UBD domains. Together with the previously reported HDAC6-UBD selective probe and its respective negative controls, SGC-UBD1031(**15**) will be a powerful tool for exploring the biological function of the USP16-UBD and investigating its potential therapeutic applications.

## EXPERIMENTAL SECTION

### COMPOUND SYNTHESIS

#### General Considerations

Unless otherwise stated, all reactions were carried out under an inert atmosphere of dry argon or nitrogen utilizing glassware that was either oven (120 °C) or flame-dried. Workups and isolation of the products was conducted on the benchtop using standard techniques. Reactions were monitored using thin-layer chromatography (TLC) on SiliaPlate™ Silica Gel 60 F254 plates. Visualization of the plates was performed under UV light (254 nm) or using KMnO4 stains. Toluene was distilled over calcium hydride, and anhydrous N,N-dimethylformamide (DMF) was purchased from Fisher Sciences and used as received. Silica gel flash column chromatography was performed on Silicycle 230-400 mesh silica gel. Mono- and multidimensional NMR characterization data were collected at 298 K on a Varian Mercury 300, Varian Mercury 400, Bruker Avance II, Agilent 500, or a Varian 600. 1H NMR spectra were internally referenced to the residual solvent peak (CDCl_3_ = 7.26 ppm, DMSO-d6 = 2.50 ppm). 13C{1H} NMR spectra were internally referenced to the solvent peak (CDCl_3_ = 77.16 ppm, DMSO-d6 = 39.52 ppm). 19F NMR chemical shifts are reported in ppm with absolute reference to 1H. NMR data are reported as follows: chemical shift (δ ppm), multiplicity (s = singlet, d = doublet, t = triplet, q = quartet, m = multiplet, br = broad), coupling constant (Hz), integration. Coupling constants have been rounded to the nearest 0.05 Hz. The NMR spectra were recorded at the NMR facility of the Department of Chemistry at the University of Toronto. Infrared spectra were recorded on a Perkin-Elmer Spectrum 100 instrument equipped with a single-bounce diamond/ZnSe ATR accessory and are reported in wavenumber (cm-1) units. High resolution mass spectra (HRMS) were obtained on a micromass 70S-250 spectrometer (EI) or an ABI/SciexQStar Mass Spectrometer (ESI) or a JEOL AccuTOF model JMS-T1000LC mass spectrometer equipped with an IONICS® Direct Analysis in Real Time (DART) ion source at the Advanced Instrumentation for Molecular Structure (AIMS) facility of the Department of Chemistry at the University of Toronto. Analytical HPLC analyses were carried out on an Agilent 1100 series instrument equipped with a Phenomenex KINETEX® column (2.6 μm, C18, 50x4.6 mm). A linear gradient starting from 5% acetonitrile and 95% water (0.1% formic acid) to 95% acetonitrile and 5% water (0.1% formic acid) over 4 minutes followed by 5 minutes of elution at 95% acetonitrile and 5% water (0.1% formic acid) was employed. Flow rate was 1 mL/min, and UV detection was set to 254 nm and 214 nm. HPLC analyses were conducted at room temperature. All compounds submitted for testing were at ≥95% purity (by HPLC, by UV detection at 254nm) unless otherwise stated. Purities of all compounds were estimated to be >95% as no significant impurities were detected in the chromatogram.

All the compounds were synthesised following the reported general procedures.^(17)^ Compounds **1, 2, 5, 8, 9,** and **10** were also reported.^(17)^

**3-(8-chloro-3-(2-((1-(2-methoxyphenyl)ethyl)amino)-2-oxoethyl)-4-oxo-3,4-dihydroquinazolin-2-yl)propanoic acid (3):** ^1^H NMR (DMSO-*d_6_*, 500 MHz): δ 8.76 (d, *J* = 8.0 Hz, 1H), 8.04 (dd, *J* = 7.9, 1.5 Hz, 1H), 7.94 (dd, *J* = 7.8, 1.4 Hz, 1H), 7.45 (t, *J* = 7.9 Hz, 1H), 7.33 (dd, *J* = 7.4, 1.8 Hz, 1H), 7.21 (ddd, *J* = 8.6, 7.5, 1.7 Hz, 1H), 6.98 – 6.89 (m, 2H), 5.22 (p, *J* = 7.1 Hz, 1H), 4.95 – 4.83 (m, 2H), 3.79 (s, 3H), 2.99 (qt, *J* = 17.5, 6.8 Hz, 2H), 2.78 (td, *J* = 6.8, 2.1 Hz, 2H), 1.32 (d, *J* = 6.9 Hz, 3H). ^13^C{^1^H} NMR (DMSO-*d_6_,* 125 MHz,): δ 173.6, 165.3, 160.7, 157.4, 155.9, 143.1, 134.5, 132.1, 130.4, 127.9, 126.9, 125.6, 125.4, 121.3, 120.4, 110.9, 55.4, 45.4, 43.3, 30.0, 29.1, 21.5. HRMS: ESI+, calcd. for C_22_H_23_ClN_3_O_5_ 444.13207 [M+H]^+^, found 444.13149.

**3-(8-chloro-3-(2-((1-(2-methoxyphenyl)ethyl)(methyl)amino)-2-oxoethyl)-4-oxo-3,4-dihydroquinazolin-2-yl)propanoic acid (4):** ^1^H NMR (CD_3_OD, 400 MHz, compound exists as a mixture of rotamers) δ 8.17 – 8.05 (m, 1H), 7.90 – 7.80 (m, 1H), 7.49 – 7.28 (m, 4H), 7.10 – 6.93 (m, 2H), 5.94 (q, J = 7.1 Hz, 0.3H), 5.72 (d, J = 17.0 Hz, 0.2H), 5.56 (q, J = 7.0 Hz, 0.5H), 5.37 (d, J = 16.9 Hz, 0.5H), 5.31 – 5.04 (m, 0.5H), 4.02 (s, 2H), 3.93 – 3.77 (m, 2H), 3.12 – 2.96 (m, 2H), 2.92 – 2.82 (m, 2H), 2.79 (s, 1H), 2.56 (s, 2H), 1.73 (d, J = 7.0 Hz, **2**H), 1.50 (d, J = 7.1 Hz, **1**H).

**3-(8-Chloro-4-oxo-3-(2-oxo-2-(((tetrahydro-2*H*-pyran-3-yl)methyl)amino)ethyl)-3,4-dihydroquinazolin-2-yl)propanoic Acid (6):** ^1^H NMR (DMSO-*d*_6_, 500 MHz): *δ* 12.18 (br s, 1H), 8.33 (t, *J* = 5.8 Hz, 1H), 8.05 (dd, *J* = 8.0, 1.4 Hz, 1H), 7.96 (dd, *J* = 7.8, 1.4 Hz, 1H), 7.47 (t, *J* = 7.9 Hz, 1H), 4.80 (s, 2H), 3.75 (ddd, *J* = 11.2, 4.0, 1.7 Hz, 1H), 3.73–3.66 (m, 1H), 3.28 (td, *J* = 10.8, 2.8 Hz, 1H), 3.09–2.92 (m, 5H), 2.79 (t, *J* = 6.7 Hz, 2H), 1.78–1.70 (m, 1H), 1.70–1.62 (m, 1H), 1.60–1.51 (m, 1H), 1.50–1.37 (m, 1H), 1.26–1.13 (m, 1H). ^13^C{^1^H} NMR (DMSO-*d*_6_, 126 MHz): *δ* 173.6, 166.3, 160.7, 157.4, 143.1, 134.5, 130.4, 126.9, 125.4, 121.4, 70.4, 67.4, 45.5, 40.9, 35.9, 29.9, 29.1, 26.9, 24.8. IR (ATR): 3289, 2935, 1662, 1598, 1561, 1442, 1389, 1219, 1169, 1088, 984, 765 cm^-1^. HRMS: ESI+, calcd. for 408.1321 for C_19_H_23_N_3_O_5_Cl, found 408.1330.

**3-(8-chloro-3-(2-((2-ethoxybenzyl)amino)-2-oxoethyl)-4-oxo-3,4-dihydroquinazolin-2-yl)propanoic acid (7):** ^1^H NMR (500 MHz, DMSO-d6): δ 12.20 (s, 1H), 8.66 (t, J = 5.8 Hz, 1H), 8.06 (dd, J = 8.0, 1.4 Hz, 1H), 7.96 (dd, J = 7.7, 1.4 Hz, 1H), 7.47 (t, J = 7.9 Hz, 1H), 7.22 (t, J = 7.3 Hz, 2H), 6.96 (d, J = 8.2 Hz, 1H), 6.91 (t, J = 7.4 Hz, 1H), 4.90 (s, 2H), 4.30 (d, J = 5.8 Hz, 2H), 4.04 (q, J = 6.9 Hz, 2H), 3.05 (t, J = 6.8 Hz, 2H), 2.81 (t, J = 6.7 Hz, 2H), 1.34 (t, J = 6.9 Hz, 3H). ^13^C{^1^H} NMR (125 MHz, DMSO-d6): δ 173.6, 166.3, 160.7, 157.3, 156.0, 143.1, 134.5, 130.5, 128.2, 127.9, 126.9, 126.4, 125.4, 121.4, 120.1, 111.4, 63.3, 45.5, 37.6, 29.8, 29.1, 14.7.8

**3-(8-Chloro-3-(2-((2-fluorobenzyl)(methyl)amino)-2-oxoethyl)-4-oxo-3,4-dihydroquinazolin-2-yl)propanoic Acid (11):** ^1^H NMR (DMSO-*d*_6_, 500 MHz, compound exists as a mixture of rotamers): *δ* 12.27 (br s, 1H), 8.08–8.02 (m, 1H), 7.97 (d, *J* = 7.7 Hz, 1H), 7.53–7.38 (m, 1.73H), 7.38–7.24 (m, 2H), 7.24–7.15 (m, 1.27H), 5.26–5.11 (m, 2H), 4.87–4.52 (m, 2H), 3.15 (s, 2H), 3.06–2.91 (m, 2H), 2.85 (s, 1H), 2.83–2.75 (m, 2H). **IR** (ATR): 1679, 1651, 1597, 1490, 1447, 1405, 1329, 1273, 1221, 1173, 1110, 761 cm^-1^. HRMS: ESI+, calcd. for 432.1121 for C_21_H_20_N_3_O_4_FCl, found 432.1126.

**(*S*)-3-(8-Chloro-3-(2-((1-(2-fluorophenyl)ethyl)amino)-2-oxoethyl)-4-oxo-3,4-dihydroquinazolin-2-yl)propanoic Acid (12):** ^1^H NMR (DMSO-*d*_6_, 500 MHz): *δ* 12.18 (br s, 1H), 8.92 (d, *J* = 7.6 Hz, 1H), 8.04 (dd, *J* = 8.0, 1.4 Hz, 1H), 7.95 (dd, *J* = 7.8, 1.4 Hz, 1H), 7.49–7.41 (m, 2H), 7.32–7.26 (m, 1H), 7.19 (td, *J* = 7.5, 1.3 Hz, 1H), 7.15 (ddd, *J* = 10.8, 8.2, 1.3 Hz, 1H), 5.16 (p, *J* = 7.1 Hz, 1H), 4.93–4.83 (m, 2H), 3.07–2.89 (m, 2H), 2.82–2.72 (m, 2H), 1.39 (d, *J* = 6.9 Hz, 3H). ^13^C{^1^H} NMR (DMSO-*d*_6_, 126 MHz): *δ* 173.5, 165.6, 160.7, 159.4 (d, *J*_C-F_ = 244.1 Hz), 157.3, 143.1, 134.5, 131.0 (d, *J*_C-F_ = 13.9 Hz), 130.4, 128.8 (d, *J*_C-F_ = 8.2 Hz), 127.2 (d, *J*_C-F_ = 4.3 Hz), 126.9, 125.4, 124.5 (d, *J*_C-F_ = 3.3 Hz), 121.3, 115.3 (d, *J*_C-F_ = 21.7 Hz), 45.3, 42.9 (d, *J*_C-F_ = 3.3 Hz), 29.8, 29.0, 21.4. IR (ATR): 1679, 1649, 1598, 1490, 1447, 1404, 1223, 1172, 760 cm^-1^. HRMS: ESI+, calcd. for 430.0964 for C_21_H_18_N_3_O_4_FCl, found 430.0971.

**(*S*)-3-(8-Chloro-3-(2-((1-(2-fluorophenyl)ethyl)(methyl)amino)-2-oxoethyl)-4-oxo-3,4-dihydroquinazolin-2-yl)propanoic Acid (13):** ^1^H NMR (CDCl_3_, 500 MHz, compound exists as a mixture of rotamers): *δ* 8.12 (d, *J* = 7.9 Hz, 1H), 7.74 (dd, *J* = 7.8, 1.4 Hz, 1H), 7.40–7.19 (m, 4H), 7.18–6.97 (m, 2H), 6.01 (q, *J* = 7.1 Hz, 0.6H), 5.47 (d, *J* = 17.4 Hz, 0.5H), 5.43 (q, *J* = 7.2 Hz, 0.4H), 5.19 (d, *J* = 16.6 Hz, 0.5H), 4.95 (d, *J* = 16.4 Hz, 0.5H), 4.74 (d, *J* = 16.6 Hz, 0.5H), 3.15– 2.99 (m, 3H), 2.98–2.87 (m, 1H), 2.85 (s, 1.37H), 2.70 (s, 1.63H), 1.73 (d, *J* = 6.9 Hz, 1.37H), 1.52 (d, *J* = 7.1 Hz, 1.63H). ESI+, calcd. for 446.1277 for C_22_H_22_N_3_O_4_FCl, found 446.1272.

**3-(8-chloro-3-(2-((1-(2,6-difluorophenyl)ethyl)amino)-2-oxoethyl)-4-oxo-3,4-dihydroquinazolin-2-yl)propanoic acid (14):** 1H NMR (500 MHz, DMSO-d6) δ 12.17 (s, 1H), 8.97 (d, J = 7.0 Hz, 1H), 8.01 (dd, J = 8.0, 1.5 Hz, 1H), 7.93 (dd, J = 7.7, 1.4 Hz, 1H), 7.44 (t, J = 7.9 Hz, 1H), 7.33 (tt, J = 8.3, 6.4 Hz, 1H), 7.04 (t, J = 8.4 Hz, 2H), 5.24 (p, J = 7.2 Hz, 1H), 4.84 (s, 2H), 2.93 (qt, J = 17.5, 6.7 Hz, 2H), 2.75 (t, J = 6.6 Hz, 2H), 1.49 (d, J = 7.2 Hz, 3H). 13C{1H} NMR (126 MHz, DMSO-d6) δ 173.5, 165.7, 161.5 (d, J = 8.7 Hz), 160.6, 159.1 (d, J = 8.6 Hz), 157.3, 143.1, 134.5, 130.4, 129.3 (t, J = 10.6 Hz), 126.8, 125.4, 121.2, 119.0 (t, J = 17.1 Hz), 111.9 (d, J = 25.3 Hz), 45.1, 29.8, 28.9, 19.7. HRMS: ESI+, calcd. for C_21_H_19_N_3_O_4_F_2_Cl 450.1027; found 450.1028

**(S)-3-(8-chloro-3-(2-((1-(2,6-difluorophenyl)ethyl)amino)-2-oxoethyl)-4-oxo-3,4-dihydroquinazolin-2-yl)propanoic acid (15):** ^1^H NMR (400 MHz, DMSO-*d*_6_) δ 12.14 (s, 1H), 8.97 (d, *J* = 7.0 Hz, 1H), 8.03 (dd, *J* = 8.0, 1.4 Hz, 1H), 7.96 (dd, *J* = 7.8, 1.4 Hz, 1H), 7.47 (t, *J* = 7.9 Hz, 1H), 7.35 (tt, *J* = 8.5, 6.5 Hz, 1H), 7.06 (t, *J* = 8.4 Hz, 2H), 5.24 (p, *J* = 7.1 Hz, 1H), 4.84 (s, 2H), 3.01 – 2.84 (m, 2H), 2.75 (t, *J* = 6.6 Hz, 2H), 1.50 (d, *J* = 7.2 Hz, 3H). ^13^C{^1^H} NMR (126 MHz, DMSO-*d*_6_) δ 173.5, 165.7, 161.5 (d, *J* = 8.6 Hz), 160.6, 159.1 (d, *J* = 8.6 Hz), 157.3, 143.0, 134.5, 130.4, 129.3 (t, *J* = 10.6 Hz), 126.9, 125.4, 121.2, 119.0 (t, *J* = 17.0 Hz), 113.7 – 110.0 (m), 45.0, 29.8, 28.9, 19.7 (t, *J* = 1.6 Hz). HRMS: ESI+, calcd. for C_21_H_19_N_3_O_4_F_2_Cl 450.1027; found 450.1027. [α]^D^ = −70° (6.8 mM in DMSO).

**(R)-3-(8-chloro-3-(2-((1-(2,6-difluorophenyl)ethyl)amino)-2-oxoethyl)-4-oxo-3,4-dihydroquinazolin-2-yl)propanoic acid (16):** ^1^H NMR (400 MHz, DMSO-*d*_6_) δ 12.14 (s, 1H), 8.96 (d, *J* = 7.0 Hz, 1H), 8.02 (dd, *J* = 7.9, 1.1 Hz, 1H), 7.95 (dd, *J* = 7.9, 1.1 Hz, 1H), 7.46 (t, *J* = 7.9 Hz, 1H), 7.40 – 7.24 (m, 1H), 7.05 (t, *J* = 8.2 Hz, 2H), 5.23 (p, *J* = 7.0 Hz, 1H), 4.83 (s, 2H), 3.07 – 2.82 (m, 2H), 2.74 (t, *J* = 6.6 Hz, 2H), 1.49 (d, *J* = 7.2 Hz, 3H). ^13^C{^1^H} NMR (101 MHz, DMSO-*d*_6_) δ 173.5, 165.7, 161.5 (d, *J* = 8.7 Hz), 160.6, 159.1 (d, *J* = 8.6 Hz), 157.3, 143.0, 134.5, 130.4, 129.3 (t, *J* = 10.5 Hz), 126.9, 125.4, 121.2, 119.0 (t, *J* = 17.1 Hz), 113.7 – 110.0 (m), 45.0, 29.8, 28.9, 19.7 (t, *J* = 1.6 Hz). HRMS: ESI+, calcd. for C_21_H_19_N_3_O_4_F_2_Cl 450.1027; found 450.1039. [α]^D^ = 71° (6.3 mM in DMSO)

## PROTEIN PRODUCTION

### EXPRESSION

All proteins were produced as described previously ^19^. Briefly, UBD domains of recombinant proteins were expressed in *E. coli* or S*f*9 cells using constructs with His₆ and/or AviTag sequences for biotinylation in pET28a-LIC, p28BIOH-LIC, pfBOH-LIC or pNICBIO2 vectors. *E. coli* strains BL21 (DE3) CodonPlus or BirA were used for HDAC6 ^1109–1213, 1109–1215^, USP13 ^183–307^, USP16 ^8–185,25-185^, USP33 ^29–134^, USP49 ^1–115^, USP5 ^171–290^, USP51 ^176–305^, and BRAP^304–390^. All cultures were grown in M9 media with 50 μM ZnSO₄, appropriate antibiotics, 0.5 mM IPTG, and 1× biotin (10 mM). Cultures were induced at OD₆₀₀ ∼0.8 and incubated overnight at 15 °C. S*f*9 cells were used to express USP16 ^1–823, 1–823 R84A^. The wild-type genes of the two USP16 constructs were PCR-amplified and subcloned into pFBOH-LIC vector. The resulting plasmids were then transformed into DH10Bac™ Competent *E. coli* (Invitrogen) cells, and recombinant viral bacmid DNA was purified. This was followed by the generation of recombinant baculovirus for baculovirus-mediated protein production in Sf9 insect cells ^21^.

### PURIFICATION

Cells were harvested at low speed (7,000-12,000 ×g, 10 minutes at 10 °C in a Beckman Coulter centrifuge) and lysed by sonication (5–10 min, Sonicator 3000, Misoni) following extraction buffer containing Tris (pH 8), NaCl, 1 mM TCEP, glycerol, benzonase, PMSF, benzamidine, and NP-40 (Sf9 only). Lysates were clarified at high speed (36,000 ×g, 60 min, 10 °C) in a Beckman Coulter centrifuge, incubated with Ni-NTA resin (Qiagen), and eluted with 250-300 mM imidazole. HDAC6(1109–1213) was cleaved with thrombin during dialysis (100 U/day, 2 mM CaCl₂, 5–7 days), followed by reverse Ni-NTA purification. HDAC6 ^1109–1213^, HDAC6 ^1109–1215^, USP13 ^183–307^, USP16 ^8–185, 25-185^, USP33 ^29–134^, USP49 ^1–115^, USP5 ^171–290^, USP51 ^176–305^, and BRAP ^304–390^ were further purified with size-exclusion chromatography (S75 16/60-) while ion exchange (IEX) chromatography MonoQ was used for USP16 ^1–823, 1–823 R84A^ purification.

All proteins were purified to 95% purity, assessed by SDS-PAGE, pooled, concentrated, snap-frozen, and stored at −80 °C. Protein identity was confirmed by LC-MS.

### ISOTHERMAL TITRATION CALORIMETRY

UPS16 ^8–185^ and HDAC6 ^1109-1213^ containing the UBDs were buffer-exchanged into PBS buffer (137 mM NaCl, 2.7 mM KCl, 10 mM Na_2_HPO_4_, and 1.8 mM KH_2_PO_4_ at pH 7.5) with 0.005% Tween-20 (v/v) and 0.75% DMSO (v/v)) using dialysis to eliminate buffer mismatches. All proteins were set to 10 µM, while the SGC-UBD1031 and SGC-UBD1031N were prepared at a concentration of 60 µM to ensure a proper molar ratio for titration. All solutions were spun down and degassed under a vacuum to remove air bubbles before loading into the ITC system (Nano ITC, TA instruments). The sample cell was loaded with 300 µL of test protein, while the reference cell was filled with degassed water. The syringe was loaded with 50 µL of the ligand solution. Titrations were conducted at 25°C with a series of 24 × 2 μL injections (with 0.167 µL per injection delivered to the sample cell) at intervals of 180s allowing baseline stabilization between injections. The stirring speed was maintained at 300 rpm throughout the experiment to ensure proper mixing. A preliminary injection of ligand 0.5 µL was performed to account for syringe-tip dilution effects but was excluded from data analysis. Raw data was then processed using Nano Analyze Software, TA Instruments. The heat change per injection was integrated and plotted against the molar ratio of ligand to protein. The binding isotherm was fitted using the independent binding model to extract thermodynamic parameters, including binding affinity (KD). Each experiment was performed in triplicate to ensure reproducibility.

### SURFACE PLASMON RESONANCE

SPR experiments were conducted using a Biacore™ 8K (Cytiva) at 20°C. Biotinylated UBD domains of USP16 ^25-185^ and HDAC6 ^1109-1215^, with approximately 1000-1200 response units (RU), were immobilized onto the flow cell two of two channels of a streptavidin-conjugated SA chip according to the manufacturer’s protocol. Flow cell one of each channel was kept empty to be used as the reference for subtraction for each channel. For the SPR run, compounds were serially titrated down (1:2) from 5 µM concentration in HBS buffer (20 mM HEPES pH 7.4, 150 mM NaCl, 0.005% Tween-20 (v/v) and 1 mM TCEP) with final DMSO maintained at 2% (v/v). Binding experiments were performed using multicycle kinetics analysis and analyte solutions were injected over the test ligand surface at a flow rate of 40 µL/min for association time, 300s, followed by a dissociation phase of 300s in the running buffer at 20°C. The dissociation constant (KD) values were determined using steady-state affinity 1:1 binding with Biacore™ Insight Evaluation software (Cytiva). All measurements were performed in triplicate to ensure reproducibility.

### UBIQUITIN RHODAMINE DUB ASSAY

The assay was conducted in the black 384-well round bottom low-volume polypropylene plates (Greiner Bio-One). SGC-UBD1031 and SGC-UBD1031N were serially titrated down (1:3) from 100 µM concentration in the assay buffer 20 mM Tris pH 7.5, 150 mM NaCl, 1 mM DTT, 0.002% TritonX-100 (v/v), with final DMSO maintained at 0.75% (v/v) containing 16 nM of either USP16^1-823^ or catalytically dead mutant USP16^1-823, R84A^. The reaction was initiated by transferring substrate Ubiquitin-rhodamine-110 (UBPBio) to a final concentration of 500 nM to the wells followed by a brief mixing by gentle pipetting (using Bravo Automated Liquid Handling Platform, Agilent) following a short spin down at 300 g (eppendorf Centrifuge 5810R). The final reaction mixture consisted of 60 µL/well. Negative control wells contained substrate and buffer without the enzyme. Fluorescence was measured continuously at excitation wavelength, 485 nm, and emission wavelength, 528 nm using the BioTek Synergy H1 microplate reader (BioTek) at room temperature for over 30 minutes. The linear portion of the plot was used for regression analysis.The data were analyzed with GraphPad Prism 9.2.0. using four-parameter dose-response curve. Each experiment was performed in triplicates to ensure reproducibility.

### NANOBRET ASSAY

HEK293T cells were plated in 6-well plates (8 × 10^5^/well) in DMEM supplemented with 10% FBS (Wisent), penicillin (100 U/mL), and streptomycin (100 μg/mL). After 4 h, cells were co-transfected with 0.02 μg of N-terminally NanoLuc-tagged HDAC6 or USP16, 0.4 μg of N-terminally HaloTag-tagged ISG15, and 1.6 μg of empty vector. The following day cells were trypsinized and seeded in a 384-well white plate (20 μl/well) in DMEM F12 (no phenol red, 4% FBS) +/– HaloTag NanoBRET 618 Ligand (1 μL/mL, Promega) and +/– compounds or DMSO. Four hours later, 5 μL/well of NanoBRET Nano-Glo substrate (8 μL/mL in DMEM no phenol red, Promega) was added, and 460 nm donor and 618 nm acceptor signals were read immediately after substrate addition using a CLARIOstar microplate reader (Mandel). Mean corrected NanoBRET ratios (mBU) were determined by subtracting the mean of 618/460 signal from cells without NanoBRET 618 Ligand ×1000 from the mean of 618/460 signal from cells with NanoBRET 618 Ligand ×1000. The IC_50_ values were determined using GraphPad Prism 10 software.

### CELL GROWTH ASSAY

HCT116 and HEK293T cells were plated in 96-well plates in RPMI or DMEM, respectively, supplemented with 10% FBS (Wisent) and penicillin (100 U/mL), and streptomycin (100 μg/mL) and treated with 30 μM compounds or DMSO control. The confluency was measured with IncuCyte™ ZOOM live cell imaging device (Essen Bioscience) and analyzed with IncuCyte™ ZOOM (2015A) software.

## Author Contributions

Compounds were tested by M.S, M.K.M, M.K.S. Compounds were designed by V.S and synthesized by B.M, J.L, A.D, J.B. Manuscript preparation was done by M.S, M.K.S, V.S, R.J.H. Guidance and advice throughout the project were provided by V.S, R.J.H, M.L, D.B, C.H.A, M.S

## ACKNOWLEDGEMENTS

The Structural Genomics Consortium is a registered charity (no: 1097737) that receives funds from Bayer AG, Boehringer Ingelheim, Bristol Myers Squibb, Genentech, Genome Canada through Ontario Genomics Institute [OGI-196], EU/EFPIA/OICR/McGill/KTH/Diamond Innovative Medicines Initiative 2 Joint Undertaking [EUbOPEN grant 875510], Janssen, Merck KGaA (aka EMD in Canada and US), Pfizer and Takeda. B.M was supported by NSERC CREATE training award and M.S. by a Mitacs postdoctoral fellowship.

