## Supplementary Information for "Development of a dual chemical probe for the USP16 and HDAC6 zinc-finger ubiquitin-binding domain"

**Table 1**

| Compound ID | Compound Structure | SPR Kd μM (USP16) | SPR Kd μM (HDAC6) |
|---|---|---|---|
| DATR31 | 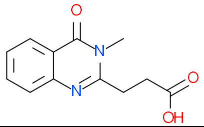 | 130 | 2.7 |
| UBXXML36 | 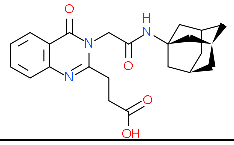 | 38 | 3.8 |
| UBXXML47 | 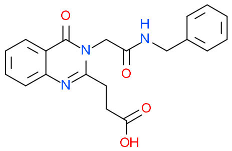 | 11 | 1.1 |
| UBXXML264 | 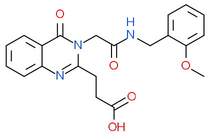 | 13 | 0.41 |
| UBXXML299 | 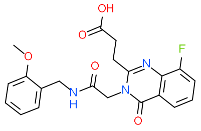 | 11 | 1.9 |
| UBXXML9098 | 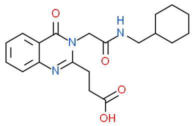 | 4.8 | 0.17 |

| | | | |
|---|---|---|---|
| UBXXML9099 | 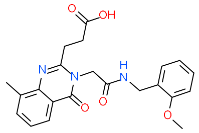 | 3 | 0.25 |
| UBXXML23602 | 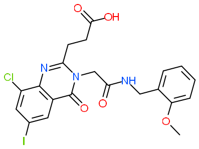 | 1.1 | 0.08 |

**Figure 1: HPLC traces of representative compounds**

**Compound 3**

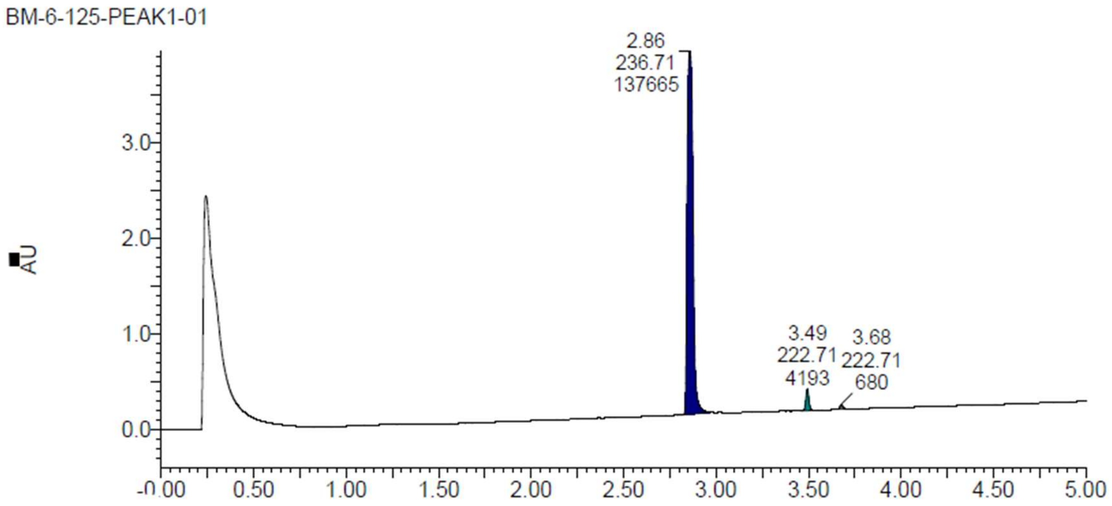


**Compound 4**

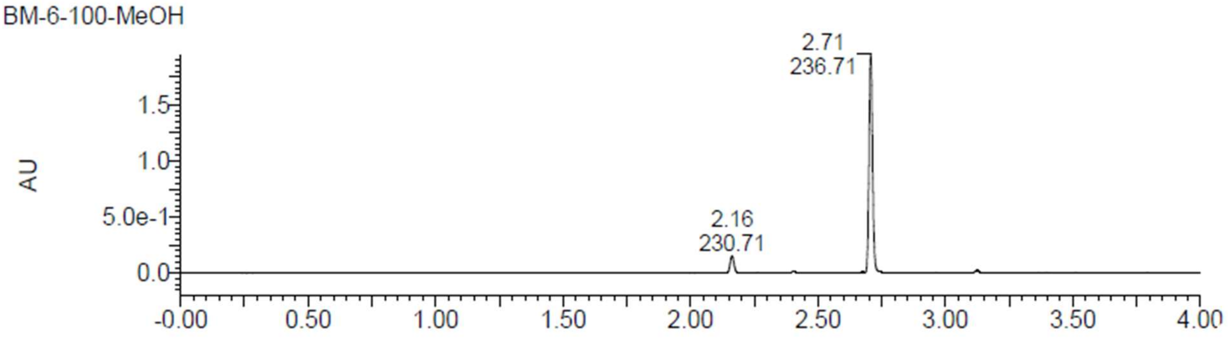

**Compound 6**

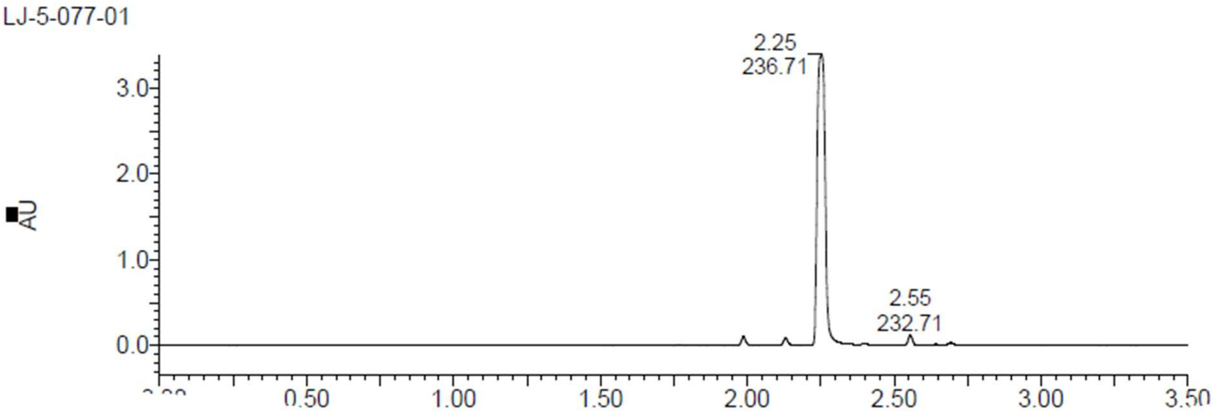


**Compound 12**

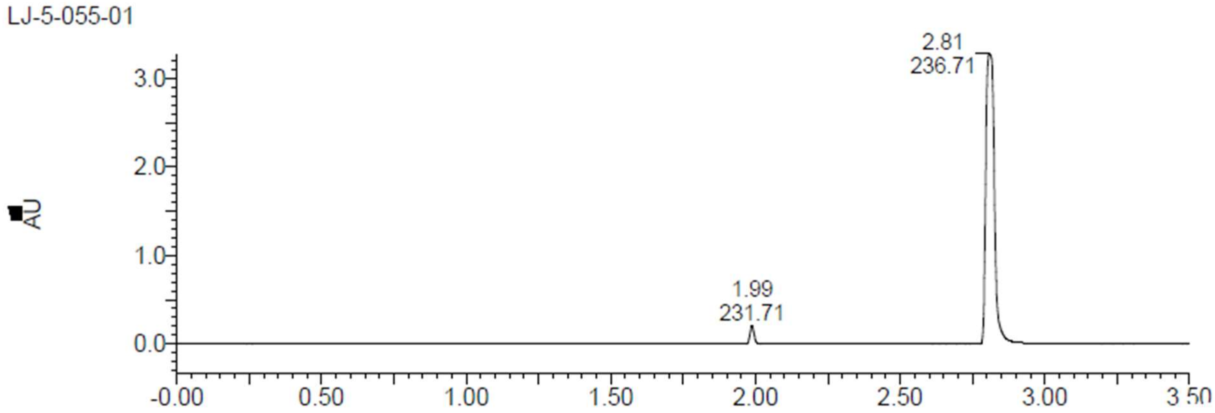


**Compound 14**

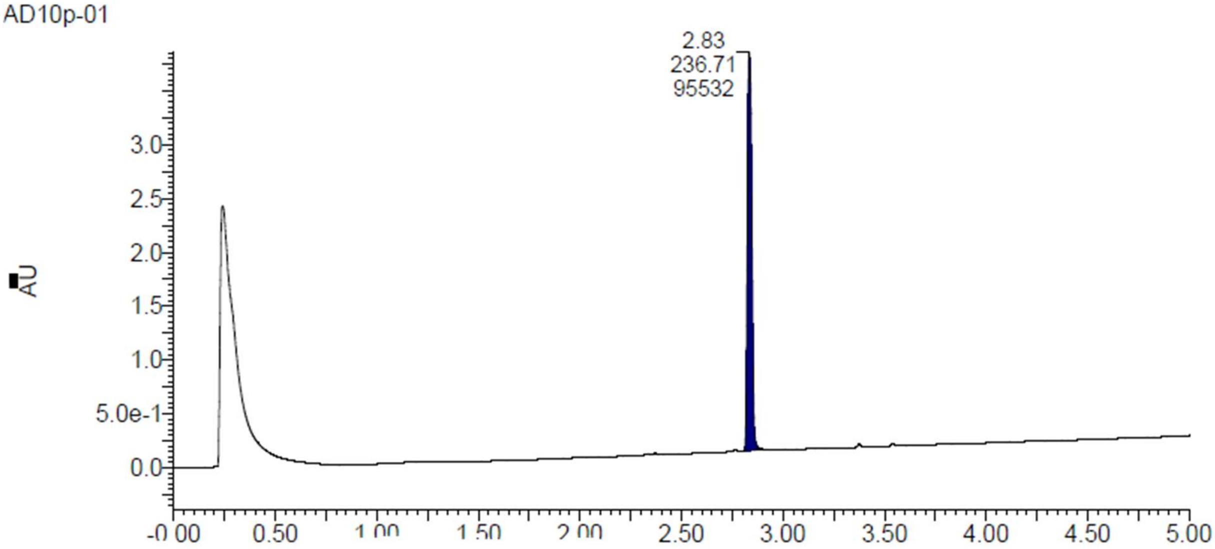

**Compound 15**

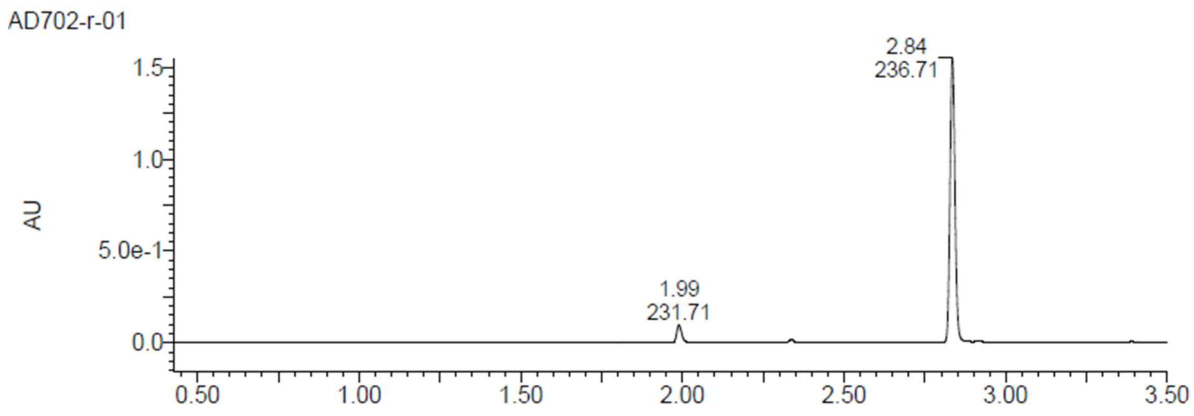


**Compound 16**

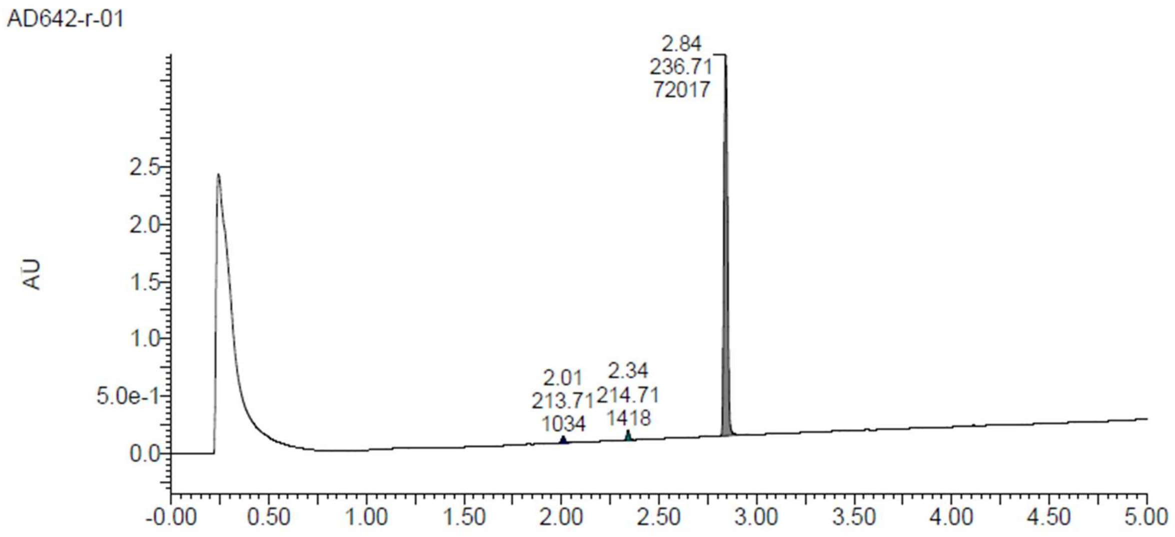
